# P53 toxicity is a hurdle to CRISPR/CAS9 screening and engineering in human pluripotent stem cells

**DOI:** 10.1101/168443

**Authors:** Robert J. Ihry, Kathleen A. Worringer, Max R. Salick, Elizabeth Frias, Daniel Ho, Kraig Theriault, Sravya Kommineni, Julie Chen, Marie Sondey, Chaoyang Ye, Ranjit Randhawa, Tripti Kulkarni, Zinger Yang, Gregory McAllister, Carsten Russ, John Reece-Hoyes, William Forrester, Gregory R. Hoffman, Ricardo Dolmetsch, Ajamete Kaykas

## Abstract

CRISPR/Cas9 has revolutionized our ability to engineer genomes and to conduct genome-wide screens in human cells. While some cell types are easily modified with Cas9, human pluripotent stem cells (hPSCs) poorly tolerate Cas9 and are difficult to engineer. Using a stable Cas9 cell line or transient delivery of ribonucleoproteins (RNPs) we achieved an average insertion or deletion efficiency greater than 80%. This high efficiency made it apparent that double strand breaks (DSBs) induced by Cas9 are toxic and kill most treated hPSCs. Cas9 toxicity creates an obstacle to the high-throughput use CRISPR/Cas9 for genome-engineering and screening in hPSCs. We demonstrated the toxic response is *tp53*-dependent and the toxic effect of *tp53* severely reduces the efficiency of precise genome-engineering in hPSCs. Our results highlight that CRISPR-based therapies derived from hPSCs should proceed with caution. Following engineering, it is critical to monitor for *tp53* function, especially in hPSCs which spontaneously acquire *tp53* mutations.

## INTRODUCTION

The bacterial-derived CRISPR/Cas9 RNA guided nuclease has been repurposed to induce user-defined double strand breaks (DSBs) in DNA (Jinek et al., 2012). This system is revolutionizing functional genomics studies, and it is now possible to conduct genetic screens in a wide range of human cells (Hart et al., 2015; Shalem et al., 2014; Wang et al., 2014). While Cas9 does not appear to induce toxicity (Ousterout et al., 2015), there are concerns about nonspecific DNA cleavage leading to off-target mutations. To address this issue, several groups have developed methods to map off-target mutations and Cas9 variants with reduced or no off-target activity (Frock et al., 2014; Kleinstiver et al., 2016; Slaymaker et al., 2015; Tsai et al., 2014; Wang et al., 2015b). In transformed cells, Cas9 is extremely efficient with minimal side effects; however, there are some cell types in which genome engineering is less efficient. Several studies have shown that gene targeting with the same reagents consistently results in five- to twentyfold lower efficiencies in human pluripotent stem cells (hPSCs) relative to other cell types (He et al., 2016; Hsu et al., 2013; Lin et al., 2014; Lombardo et al., 2007; Mali et al., 2013). The exact cause of this reduced efficiency remains unclear but it presents a significant challenge for approaches such as genome-wide screens and for *ex vivo* therapeutic editing in hPSCs.

hPSCs derived from preimplantation embryos or by cellular reprogramming hold great promise for both genetic screening and therapeutic applications. hPSCs are genetically intact, expandable and can be differentiated into a wide variety of cell-types which are difficult to obtain from human patients (Avior et al., 2016). Despite these advantages, several challenges remain in developing a practical system for high-throughput genetic engineering of hPSCs. In addition to requiring daily feeding and expensive media, hPSCs are recalcitrant to genome modification, making techniques commonly used in other cell types and organisms difficult to implement (Hockemeyer and Jaenisch, 2016; Hockemeyer et al., 2009; Liu and Rao, 2011; Merkle et al., 2015; Song et al., 2010; Zwaka and Thomson, 2003). Enhancing the genetic toolkit in hPSCs is necessary to utilize their full potential in genetic screening, disease modelling and cell therapy. We optimized a stable system using a drug inducible Cas9 that achieves a 90% editing efficiency and determined that DSBs induced by Cas9 are toxic to hPSCs. A key finding is that DSB toxicity is the primary reason why transient CRISPR/Cas9 engineering is inefficient in hPSCs. We found that transient TP53 inhibition minimizes toxicity, leading to over a fifteen-fold increase in transgene insertion. These findings provide an explanation for the longstanding observation that hPSCs have reduced genome-engineering efficiencies and have identified that DSB-induced toxicity is a barrier to high-throughput genome-engineering in hPSCs. Our observation highlights that therapeutic use of CRISPR should proceed with caution and *tp53* activity be fully monitored after editing. This is especially important in hPSC where there is a low level of spontaneous dominant negative *tp53* mutations.

## RESULTS

### Efficient Cas9 gene disruption is toxic to hPSCs

We improved the 2-component Cas9 system developed by Gonzalez et. al., 2014., by consolidating it into a single all-in-one AAVS1 safe-harbor targeting vector with the 3^rd^ generation doxycycline (dox) inducible system and an insulator to further prevent leaky expression (henceforth iCas9; Fig. 1A, S1A). The stable Cas9 line used for this study had a normal karyotype, strong induction of Cas9 only in the presence of dox, and was properly targeted (Fig. S1B-E). The sgRNAs were delivered by lentiviruses (lentiCRISPRs). iCas9 cells were infected with 47 lentiCRISPRs targeting 16 genes and treated with dox for 8 days in a 96-well plate. DNA was then isolated and next generation sequencing (NGS) was used to quantify control and mutant allele frequencies. NGS analysis of infected cells revealed high percentages of indels (Fig. 1B). The average editing for the 47 sgRNAs was over 90% and picking 3 sgRNAs per gene identified at least 1 sgRNA that generated over 80% loss-of-function alleles (Fig. 1C). Despite efficient indel generation, it was evident only a small fraction of the hPSCs were surviving. CRISPR/Cas9 activity caused a sharp decrease in the cell number, with delayed doubling times and the presence of cellular debris. This toxicity created large variability across the wells presenting a challenge for high-throughput screens using density-dependent differentiation protocols (Table S1).

**Figure 1.**
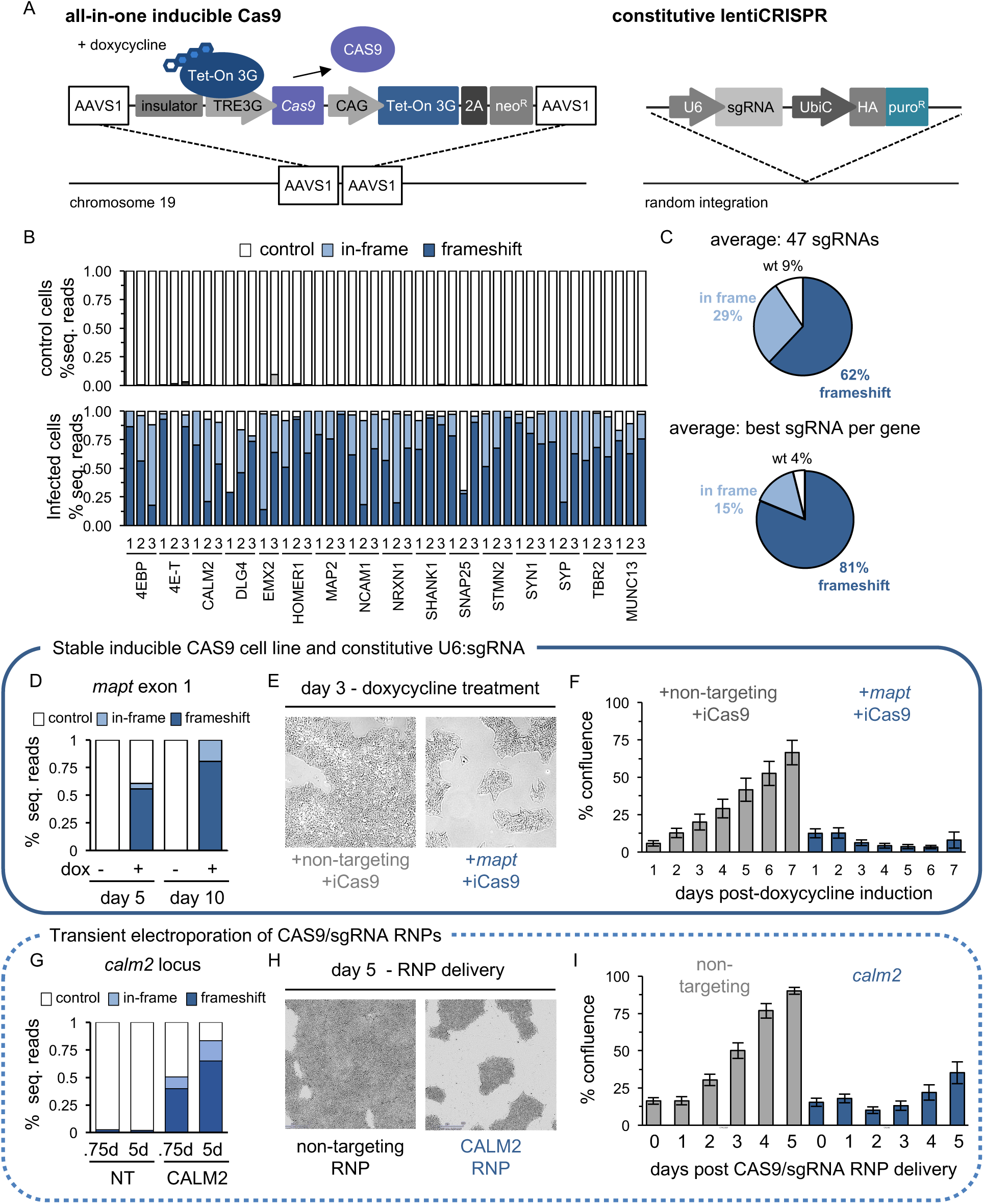
Efficient Cas9 gene disruption is toxic to hPSCs. (A) 2-component Cas9 system depicting all-in-one inducible Cas9 construct and lentiviral delivery of constitutive sgRNA. (B-D, G) NGS quantification of indels. Control reads are represented by white bars, in-frame mutations by light blue bars and frameshift mutations by dark blue bars. (B) iCas9 control cells (top) and cells infected with 47 sgRNAs (bottom) grown in the presence of dox for 8 days. >10K reads for each pooled sample (n=1) (C) Summary of efficiency and indel types generated by 47 sgRNAs. Averages shown for all 47 sgRNAs and the best sgRNA per gene. (D) Indel quantification at *mapt* locus. After 10 days of dox treatment *mapt* locus is completely edited. In the absence of dox, no editing was observed showing Cas9 is tightly controlled. >200K reads for each sample (n=1) (E) *mapt* targeting sgRNA reduces colony size relative to a non-targeting control. Bright-field image of live iCas9 cells cultured with dox for 3 days in the presence of a non-targeting or *mapt* sgRNA. (F) Quantification of toxic response to Cas9-induced DSBs in live cells. Percent confluence was measured each day in cells expressing a non-targeting or *mapt* sgRNA grown in dox. Bars represent mean. Error bars +/− 1 std. dev images from n=88 and n=96 wells respectively. The toxic response has been replicated >3 times (G) NGS quantification of indels at *calm2* locus 18 hours and 5 days (d) after electroporation of Cas9/sgRNA RNP complexes. Average indels from three independent electroporations. (H) Bright-field images of Cas9 RNP treated cells 5 days after electroporation. Electroporation of active Cas9 RNPs is toxic and decreases cell density. (I) Electroporation of Cas9 and *calm2* sgRNA containing RNPs are toxic relative to non-targeting control RNPs. Y-axis is % confluence. X-axis represents days after electroporation. Bars represent mean. Error bars +/− 1 std. dev images from 121 images per well of a 6-well plate from 3 independent electroporation (n=3). The toxic Cas9 RNP response has been replicated twice.

To study the basis of toxicity in more detail, we used the iCas9 line and a lentiCRISPR targeting *mapt*, a neuronal gene not expressed or required for survival in hPSCs. Ten days of dox treatment completely edited the *mapt* locus (Fig. 1D) and reduced colony size relative to nontargeting controls without a DSB (Fig. 1E). To quantify this, confluency was measured in live cells expressing either a non-targeting or a *mapt* sgRNA in the presence of dox (Fig. 1F). While cells expressing non-targeting controls increased confluency at a steady rate, those expressing a *mapt* sgRNA decreased confluency despite being seeded at a similar density. Despite the toxic response, *mapt* edited cells retained expression of the pluripotency markers TRA-1-60, OCT4 and SOX2 (Fig. S1). To determine if toxicity was related to off-target DSBs, we assayed the top 6 off-target sites by NGS identified by the CRISPR design tool (Hsu et al., 2013) and detected no off-target mutations (Fig. S2A and Table S2). Transient exposure to Cas9 and *calm2* targeting sgRNAs by electroporating Cas9 and sgRNA containing ribonucleoprotein (RNP) complexes also triggered a toxic response (>80% indels, Fig. 1G-I). The transient nature of RNP delivery minimizes off-target cutting (Liang et al., 2015) and further supports that DSBs at a single locus are sufficient to cause toxicity in hPSCs. We also generated H1-hESCs and 8402-iPSC lines with a dox inducible enhanced Cas9 (ieCas9) variant that greatly reduces non-specific DSBs (Fig. S1., Wells et al., 2016; Slaymaker et al., 2015). The presence of enhanced Cas9 with additional sgRNAs targeting the neuronal genes, *calm2* and *emx2*, in both hESC and iPSCs backgrounds caused a toxic phenotypic response (Fig. S2B, S2C). Cumulatively this suggests toxicity is not due to effects on other genes or many DSBs and implies that editing at a single locus is toxic.

### CRISPR screens identify an hPSC-specific toxic response to Cas9-induced DSBs

To globally test if targeting sgRNAs are toxic we conducted a large-scale pooled CRISPR screen. A total of 200 million H1-hESCs were infected at .5 MOI which is sufficient for a genome-wide screen in transformed cells. However, to control for toxicity we screened a focused 13K library at high coverage (1000x per sgRNA) across four independent conditions (Fig. 2A). 72 sgRNAs were non-targeting and the remaining targeted ~2.6K genes (5 sgRNAs/gene). All four conditions were infected with the sgRNA library with two replicates. Two conditions were grown in the absence of Cas9; the parental H1 cells and H1-iCas9 cells (−dox). To further validate the toxicity in hPSCs we generated a second inducible Cas9 based on the Shield1-destabilizing domain (DD) system (Banaszynski et al., 2006). H1s were generated with Cas9 fused to a DD tag (ddCas9) which is stabilized in the presence of Shield1 and degraded in its absence (Fig. S1). The remaining two conditions were grown in the presence of Cas9 induced by dox or Shield1, respectively. Cells were dissociated to seed new flasks and to be pelleted for DNA isolation every 4 days at 1000x sgRNA representation. Cell counts at day 4 demonstrated that iCas9 or ddCas9 hPSCs cultured with dox or Shield1 had little growth compared to H1 and iCas9 hPSCs infected with the same library, seeded at the same density, but in the absence of Cas9 induction (Fig 2B). This was reproducible and exposing the uninduced H1-iCas9 pool of infected cells to dox after passaging severely reduced cell counts relative to untreated controls (Fig. S3). NGS was used to recover spacer sequences which act as molecular barcodes to count sgRNA infected cells. All but one of 24 samples recovered 98% of expected spacer sequences, demonstrating adequate representation was maintained for most sgRNAs. Fold change was calculated for each spacer sequence by dividing each condition by the sequenced lentiviral pool (prior to infection).

**Figure 2.**
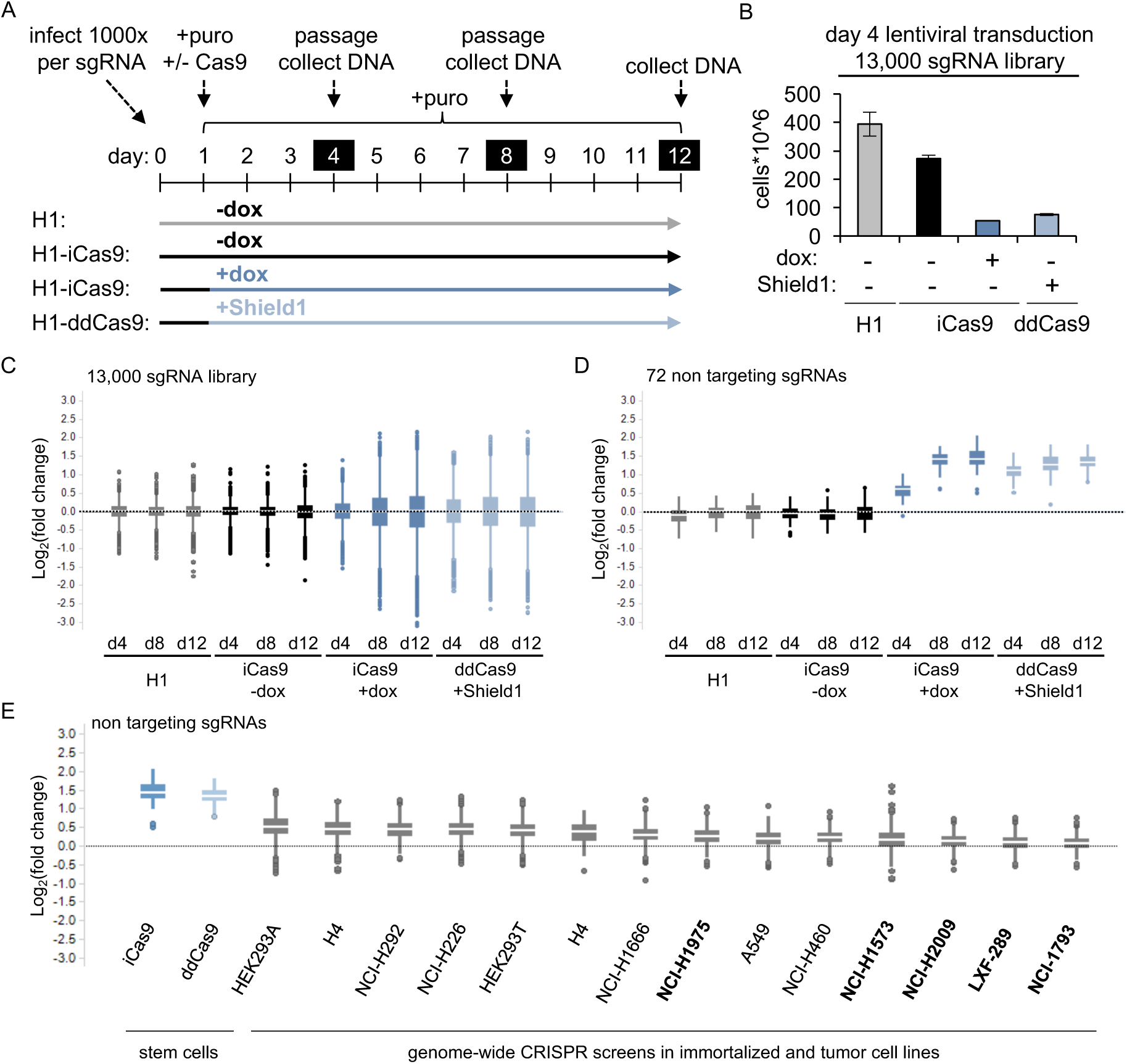
CRISPR screens identify hPSC-specific toxic response to Cas9-induced DSBs. (A) Experimental paradigm for pooled screen in hPSCs testing 13K sgRNAs in 4 independent cell lines H1: Parental depicted in light gray, H1-iCas9 minus dox depicted in black, H1-iCas9 plus dox depicted in blue, and H1-ddCas9 plus Shield1 depicted in light blue. 2000x cells for each condition were infected with each sgRNA (.5 MOI, 2.6*10^7 cells). 24 hours after lentiviral infection non-transduced cells were killed by puromycin. On days 4, 8, and 12 cells were dissociated, then counted to maintain 1000x representation for both DNA isolation and passaging. (B) Cell counts at day 4 were reduced in Cas9 positive cells (plus dox or Shield1) relative to cells grown in the absence of Cas9 (H1, iCas9 minus dox). Bars represent mean. Error bars +/− 1 std. dev from n=2 per condition. Replicate results in figure S3 (C-E) Barcode counting of genome integrated sgRNAs via NGS to measure representation of each sgRNA. (C-E) Y-axis box plots depicting log2(fold change) calculated for each sgRNA normalized to the initial 13K sgRNA library. For each box blot the median is depicted by white line flanked by a rectangle spanning Q1-Q3. n=2 per condition (C-D) X-axis plots each condition over time. (C) Fold change for the entire 13K sgRNA library. In the absence of Cas9 (grey and black) sgRNA representation does not change. In the presence of Cas9 sgRNAs both increase and decrease representation in a time-dependent manner. (D) Fold change for 72 non-targeting control sgRNAs. In the presence of Cas9, nontargeting sgRNAs enrich their representation relative to the starting library. (E) hPSCs are sensitive to DSBs. X-axis plots CRISPR screens conducted in 2 hPSC lines (Fig. 2D-day 12) and 14 additional transformed lines. In hPSCs non-targeting controls have a strong proliferative advantage over toxic DSB-inducing sgRNAs and thereby increase representation throughout the course of a CRISPR screen. The response is reduced in transformed cell lines. Lines with *tp53* mutations in bold.

Over the 12-day experiment, most sgRNAs remained distributed within +/− 1 log_2_(fold change) in uninduced conditions (Fig. 2C, File S1). In contrast, the Cas9-induced conditions displayed a time-dependent change in sgRNA representation which increased the spread of the distribution. Plotting only the non-targeting controls identified a 1.3- to 1.4-fold enrichment specific to the Cas9-induced conditions (Fig. 2D). This indicates that sgRNAs targeting the genome are globally depleted compared to non-targeting control and demonstrates that CAS9 is toxicity is widespread and over a larger number of gRNAs. To determine if this toxic response is specific to hPSCs, we evaluated the non-targeting controls across pooled CRISPR screens in other cell lines to quantify sensitivity to DSBs. Fold change was calculated for non-targeting sgRNAs from 14 additional transformed lines using a genome-wide sgRNA library. Comparing the non-targeting controls from the Cas9-induced conditions with the transformed lines demonstrated a heightened sensitivity to DSBs in hPSCs (Fig. 2E). hPSCs have a greater than 1.3-fold change while transformed cell lines show little enrichment (.05- to .51-fold change). Lastly, we exploited design flaws affecting a subset of the sgRNA library to identify additional evidence for DSB-toxicity. SNPs present in the H1-hESC genome disrupted 249 of the sgRNAs, reducing their ability to create DSBs and causing them to significantly enrich when compared to uninduced or Cas9-free parental lines (Fig. S4A). Multiple perfect cut sites were identified for 151 of the sgRNAs, which enhanced their depletion (Fig. S4B). Cumulatively, these results demonstrate hPSCs are extremely sensitive to DSBs and the effect is widespread over many sgRNAs. This toxic effect presents a significant challenge for both engineering and screening efforts.

### Cas9 DSB-induced transcriptional response in hPSCs

To further investigate the mechanism by which Cas9 causes toxicity in hPSCs, RNA-seq and differential expression analysis was performed on iCas9 cells expressing either a non-targeting or *mapt* sgRNA grown in dox for 2 days (Fig. 3A, File S2). Despite this toxic response to DSBs, the expression of pluripotency markers *oct4*, *nanog* and *sox2* were unchanged relative to nontargeting controls. However, a significant number of genes were induced in cells with a DSB relative to controls. Gene ontology analysis of the top 100 hits identified 25 genes with roles in programmed cell death (STRING-db, FDR 1.92E-08) including components of the intrinsic and extrinsic death pathways such as *bax*, *bbc3*, *fas*, and *tnfrsf10b*. Consistent with this, immunofluorescent staining of DSB induced iCas9 cells identified increases in DNA damage and apoptotic markers; phospho-histone H2A.X (pH2A.X), cleaved PARP (cPARP) and cleaved caspase-3 (CC3) (Fig. 3B, S5).

**Figure 3.**
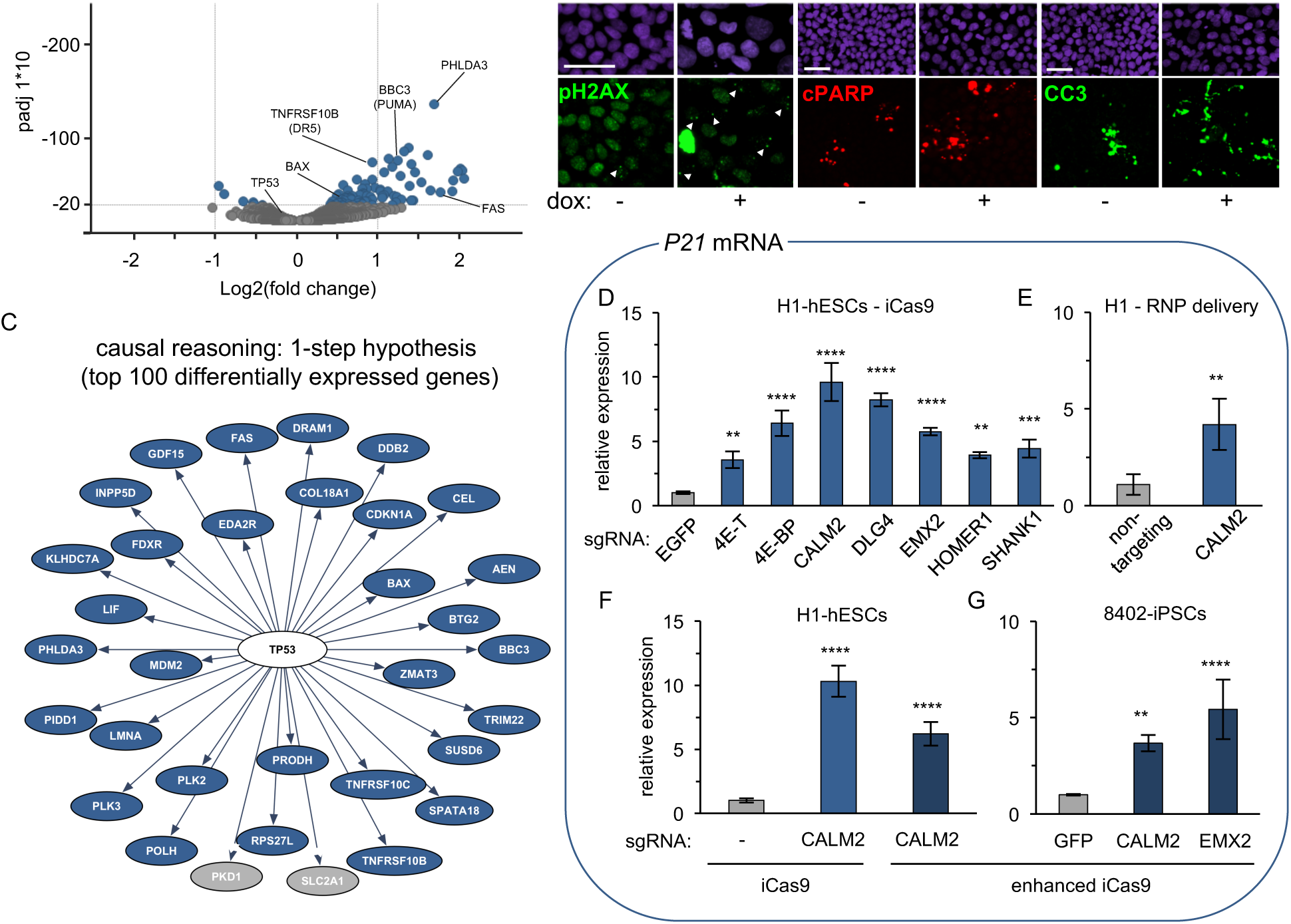
Cas9 DSB-induced transcriptional response. (A) Volcano plot depicts differential expression from RNA-seq calculated by comparing a *mapt* and non-targeting control sgRNA (n=3 per condition). Adjusted p-value (padj) on y-axis, log2(fold change) on x-axis. Each condition was cultured in the presence of dox for two days. Circles represent differentially expressed genes (blue circles padj < 1.2E-17, gray circles padj >1.2E-17). (B) High content image analysis of *mapt* sgRNA infected H1-iCas9 cells cultured with or without dox. The frequency of nuclei containing pH2AX foci increases in the presence of a DSB (+dox), p<0.001 via multiple Welch’s unpaired two-tailed t-test, n=8 wells per condition. pH2AX is green in columns 1 and 2. White triangles indicate nuclei with foci. Levels of the apoptotic marker cPARP increase in cells with a DSB (+dox), p<.001 via Welch’s unpaired two-tailed t-test, n=8 wells per condition. cPARP is red in columns 3 and 4. Immunostaining in green for cleaved caspase 3 (CC3), identified an increased area positive for CC3 debris within the dox treated iCas9 colonies, p<.05 via Welch’s unpaired two-tailed t-test, n=8 wells per condition. DAPI stained nuclei are purple Scalebar = 100um. (C) Interactome analysis identifies *tp53*-dependent changes in expression caused by Cas9-induced DSBs. The 1 -step *tp53* hypothesis accurately explains gene expression changes for 33 out of 100 differentially expressed genes. Upregulated genes in blue and downregulated genes in gray. (D-G) qPCR detects an increase of *p21* mRNA in cells treated with DSB-inducing Cas9. Y-axis is relative expression and each bar represents mean relative expression. X-axis is each sgRNA. n=3 independent mRNA samples per sgRNA, error bars +/− 1 std. dev. One-way ANOVA, equal variances **p<0.01, ***p<0.001, ****p<0.0001. (D) *p21* mRNA is induced by 7 independent sgRNAs in iCas9 cells 2 days after dox treatment. Relative expression is calculated by comparing the non-targeting control (EGFP) to each targeting sgRNA. (E) Quantification of *p21* mRNA induction 2 days following Cas9 RNP delivery. *p21* mRNA expression was measured from cells treated with non-targeting and *calm2* sgRNA. Relative expression (Y-axis) is calculated by comparing each sample to H1 hESCs electroporated with Cas9 and non-targeting sgRNA RNPs (F-G) Enhanced (e)Cas9 induces *p21* mRNA. (F) H1-hESCs with an iCas9 or an enhanced iCas9 transgene and *calm2* sgRNA have increased *p21* mRNA two days after dox treatment. Relative expression is calculated by comparing each sample to H1-iCas9 cells expressing Cas9 without an sgRNA. (G) 8402 iPSCs with an enhanced iCas9 transgene have increased *p21* mRNA two days after dox treatment in the presence of *calm2* or *emx2* targeting sgRNAs. Relative expression is calculated by comparing the non-targeting control (EGFP) to each targeting sgRNA. enhanced iCas9 (dox inducible - enhanced Streptococcus Pyogenes Cas9 1.1 (eSpyCas9(1.1)). H1-hESCs, H1 human embryonic stem cells, 8402, 8402-iPSCs human induced pluripotent stem cells.

To identify key pathways involved, an *in*-*silico* interactome analysis was performed on the top 100 differentially expressed genes (adjusted p-value cutoff of 1.2E-17). Causal reasoning algorithms consistently identified *tp53* as one of the top ranking hypotheses along with *myc*, *sp1* and *ep300* (Chindelevitch et al., 2012; Jaeger et al., 2014). These hypotheses are tightly interconnected and further investigation was focused on *tp53* because of its well-established role in the DDR (Lane, 1992). The 1-step *tp53* hypothesis accurately explained 33 out of the 100 input genes (Fig. 3C) and was consistent with *p21*, a canonical TP53 target, being the most differentially expressed gene (El-Deiry et al., 1993). In addition, 5 of 14 transformed lines had mutations in *tp53* and had reduced Cas9 induced toxicity (Fig 2E, bold). Consistent with studies demonstrating that *tp53* activity and expression are regulated post-transcriptionally we did not observe a change in *tp53* mRNA (Canman et al., 1998; Vassilev et al., 2004).

The most differentially expressed gene was *p21 (cdkn1a)* (6.12-fold, 6.6E-298 padj), a cell cycle regulator and *tp53* target with known roles in DNA damage response (DDR) (Cazzalini et al., 2010). To confirm these results, iCas9 cells were infected with 7 independent sgRNAs, treated with dox for 2 days and *p21* mRNA was measured by qPCR (Fig. 3D). The expression of *p21* was increased between 3- and 10-fold in the targeting sgRNAs compared to a non-targeting EGFP control sgRNA. Transient exposure from electroporating Cas9 and sgRNA containing ribonucleoprotein (RNP) complexes triggered a toxic response and increased *p21* expression (Fig. 3E). Additionally, the use of the enhanced Cas9 did not abrogate the induction of *p21* mRNA during DSB induction in hESCs or iPSCs, which is consistent with the toxic phenotype (Fig. 3F-G, Fig S2B-C). Both enhanced Cas9 and transient Cas9 RNP delivery minimizes off-target cutting (Liang et al., 2015, Slaymaker et al., 2015) and further supports that DSBs at a single locus are sufficient to cause toxic molecular response in hPSCs.

### Cas9 induced toxicity is *tp53*-dependent in hPSCs

To provide experimental evidence that *tp53* is functionally involved, we knocked out *tp53* in iCas9 cells by transiently transfecting 3 chemically synthesized crRNA/tracrRNA pairs targeting the *tp53* locus (Fig. 4A). The resulting mutant pool had a mixture of mutations at 3 independent sites within the *tp53* open-reading frame (ORF) (Fig. S6A). The control and *tp53* mutant pool were then infected with a *mapt* sgRNA and grown +/− dox for up to 6 days (Fig. 4B). To confirm that the transcriptional response is *tp53*-dependent, mRNA was isolated and quantified using qPCR. At day 2, control cells exhibited a strong induction of *p21* and *fas* mRNA that was significantly reduced in the *tp53* mixed mutant pool (Fig. 4C). We confirmed the involvement of TP53 and P21 proteins by measuring expression using immunofluorescence and high-content imaging. Both TP53 and P21 increased in DSB-induced controls and this is significantly reduced in the *tp53* mutant pool (Fig. 4D, S6B). Finally, the toxic response was quantified by measuring confluency during editing in the control and *tp53* mutant pool. Dox treated controls had a strong toxic response while the *tp53* mutant pool continued to grow despite DSB induction (Fig. 3E, S6C). Collectively these results demonstrate that *tp53* is required for the toxic response to DSBs induced by Cas9.

**Figure 4.**
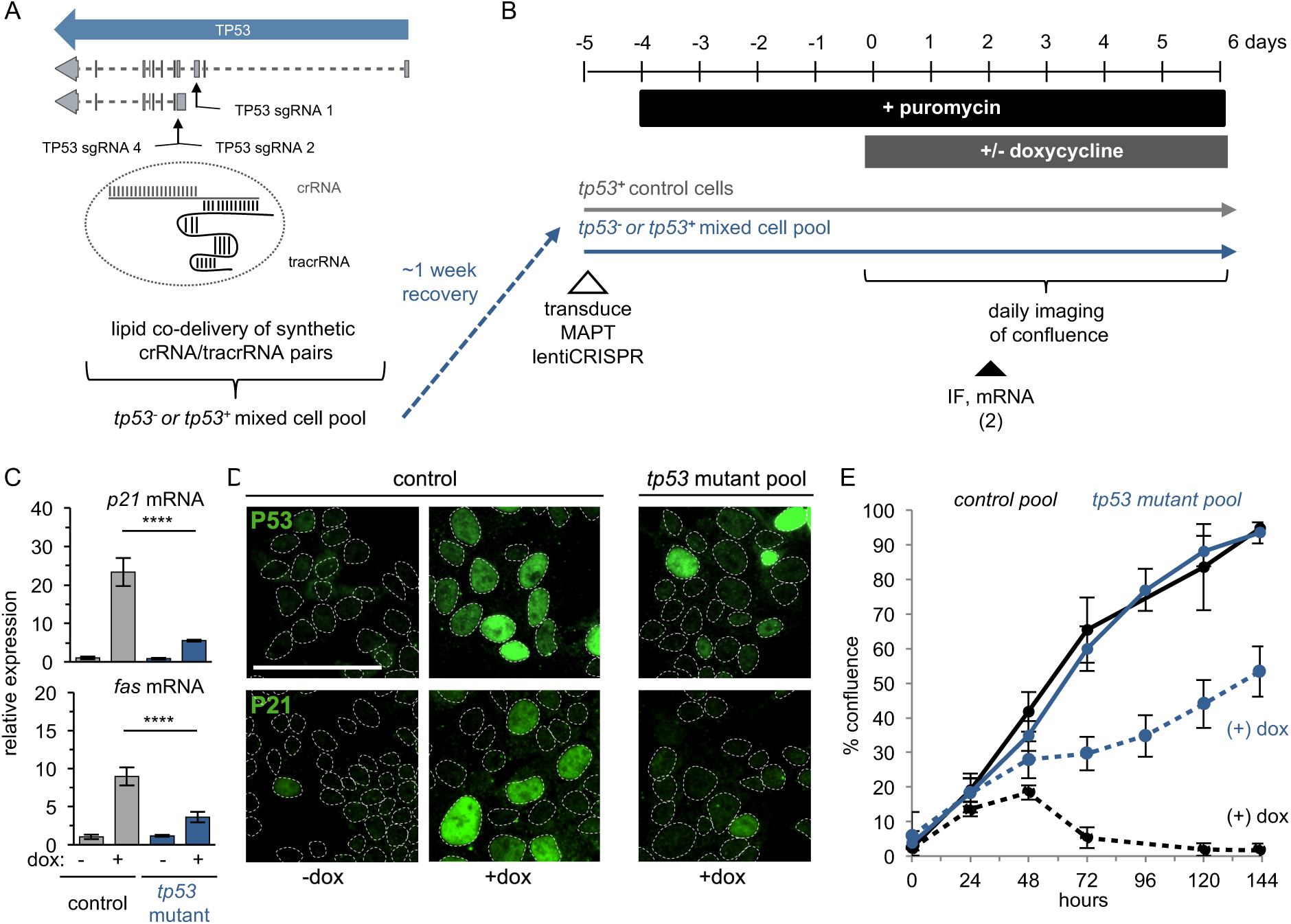
Cas9-induced toxicity is *tp53*-dependent in hPSCs. (A) Diagram showing locations of 3 synthetic crRNAs targeting the *tp53* locus. (B) Experimental paradigm for *tp53* mutant analysis. After recovering from mutagenesis, the *tp53* mutant pool and controls with an intact *tp53* were infected with the *mapt* lentiCRISPR. At the onset of the experiment, control and mutant pools were dissociated and plated into media plus or minus dox. (C) qPCR detects an induction of *p21* and *fas* mRNA in dox treated controls expressing the *mapt* sgRNA, in contrast *p21* and *fas* are significantly reduced in the *tp53* mutant pool. Relative expression (Y-axis) is calculated by comparing to untreated control cells. Each bar is mean relative expression. Genotype and dox treatment labeled on X-axis. n=3 independent mRNA samples per condition, error bars +/− 1 std. dev. One-way ANOVA, equal variances ***p<.001 ****p<0.0001. (D) In control *mapt* sgRNA infected cells, immunofluorescence staining detects DSB-dependent (+dox) increases in TP53 and P21 protein. In the *tp53* mutant pool, the percent of TP53+ and P21+ nuclei are significantly decreased via one-way ANOVA, similar variances, p<0.0001. TP53 and P21 are shown in green. DAPI co-stained nuclei are outlined in white. Scalebar = 100um (E) Cas9-induced toxic response is *tp53*-dependent. Live imaging of confluence in *mapt* sgRNA expressing iCas9 cells +/− dox in control or *tp53* mutant pool. Unlike dox treated control cells the *tp53* mutant pool continues to grow despite the induction of DSBs. Black lines indicate control and blue lines indicate tp53 mutant pool. Solid lines are without dox and dashed lines are cultured with dox. Colored circles represent mean confluency. error bars +/− 1 std.

Inserting a transgene into a specific locus by using Cas9 to stimulate HDR is a challenging task in hPSCs. We hypothesized that DNA damage-induced toxicity is in direct opposition of engineering efforts. To determine if TP53 inhibits precision engineering, we developed an assay to measure precise targeting of a transgene into the *oct4/pou5f1* locus. We used a pair of dual nickases (Ran et al., 2013) flanking the stop codon to trigger a DSB and initiate HDR with a gene trapping plasmid (Fig. S7). The gene trap does not contain a promoter or nuclear localization signal of its own and only correctly targeted cells will express a nuclear tdTomato and gain resistance to puromycin. TP53 signaling was transiently blocked by overexpressing a dominant negative p53DD transgene that inhibits the *tp53* DSB response and has been routinely used to increase reprogramming efficiency of iPSCs without causing major genome instability (Hagiyama et al., 1999; Hong et al., 2009; Schlaeger et al., 2015). The Cas9^D10A^-sgRNA(s) and *pou5f1* gene trapping plasmids were co-electroporated with or without the p53DD plasmid and scored for the number of puromycin–resistant colonies expressing nuclear tdTomato (Fig. S7B-D). TP53 inhibition greatly increased the number and size of TRA-1-60 positive colonies surviving the engineering and selection process in both 8402-iPSCs and H1-hESCs (Fig. S7B). Multiple independent experiments showed that control 8402-iPSCs and H1-hESCs had an average of 26.3 and 54.5 colonies and that p53DD significantly boosted this average to 500 and 892, respectively (Fig. S7C). TP53 inhibition resulted in a 19-fold increase in successful insertions for 8402-iPSCs and a 16-fold increase for H1-hESCs dramatically improving the efficiency of genome engineering in hPSCs.

## DISCUSSION

Genome engineering of hPSCs using Cas9 is revolutionary. However, to exploit this fully we need to increase editing efficiency and reduce toxicity. We developed a highly efficient Cas9 system in hPSCs that will be useful for genetic screening and for making collections of engineered cells. We found that DSBs induced by Cas9 triggered a *tp53*-dependent toxic response and that toxicity reduces the efficiency of engineering by at least an order of magnitude.

Recently, several groups have demonstrated that multiple cuts induced by Cas9 can cause death in transformed cells (Aguirre et al., 2016; Hart et al., 2015; Munoz et al., 2016; Wang et al., 2015a). In contrast, targeting a single locus is sufficient to kill the majority of hPSCs. Given their biological similarity to the early embryo, it is fitting that hPSCs are intolerant of DNA damage. The extreme sensitivity to DSBs may serve as a control mechanism to prevent aberrant cells from developing within an organism (Dumitru et al., 2012; Liu et al., 2013). The heightened *tp53*-dependent toxic response provides an explanation for the long-standing observation that hPSCs have inefficient rates of genome engineering. Several studies comparing indel and HDR efficiencies across cell lines identified a 5- to 20-fold reduction in hPSCs relative to transformed lines (He et al., 2016; Hsu et al., 2013; Lin et al., 2014; Lombardo et al., 2007; Mali et al., 2013). These results agree with our observation that TP53 inhibits HDR efficiency by an average of 17fold in hPSCs. While long-term TP53 inhibition can lead to increased mutational burden (Hanel and Moll, 2012), transient inhibition is well tolerated in hPSCs (Schlaeger et al., 2015; Qin et al., 2007; Song et al., 2010). TP53 inhibition may facilitate the generation of large collections of engineered hPSCs by increasing efficiency and reducing variable yields

The toxic response to Cas9 activity has important implications for gene therapy. Our observations suggest that *in vivo* genome editing in primary cells with a heightened DDR could result in significant toxicity and tissue damage. TP53 inhibition could alleviate toxicity but has the potential to increase off-target mutations and poses a risk for cancer. For *ex vivo* engineering, Cas9 toxicity combined with clonal expansion could potentially select for *tp53* mutant cells more tolerant of DNA damage. Although the mutation rate of *tp53* remains to be determined for other clinically relevant cell types, this is a serious concern for hPSCs. The basal *tp53* mutation rate in hESCs is significant and Merkle et al., 2017 have identified that 3.5 % of independent hESC lines and up to 29% of hESCs commonly used in RNA-seq databases have *tp53* mutations. Before engineering patient cells, the risks and benefits must be fully evaluated. It will be imperative to determine the spontaneous mutation rate of *tp53* in engineered cells as well as the mutational burden associated with transient *tp53* inhibition. As gene and cell therapies become tangible, it will be critical to ensure patient cells have a functional
*tp53* before and after engineering.

## EXPERIMENTAL PROCEDURES

### Lentiviral and lipid delivery of sgRNAs for Cas9 mutagenesis

During replating lentiCRISPRs were added to a single cell suspension of 2*10^5^ cells in 2ml E8 (STEMCELL TECH.-05940) with .8uM thiazovivin (Selleckchem-S1459). After 24 hours, cells were maintained in 2ug/ml to puromycin (Corning-61-385-RA). At the onset of each mutagenesis experiment Shield1 (Clontech-631037) at .5uM and dox (Clontech-631311) at 2 ug/ml were added to induce Cas9. To disrupt *tp53*, 3 *tp53*-targeting crRNAs each at 30nM were delivered with 90nM tracrRNA (IDT). RNAimax (ThermoFisher-13778150) was used to deliver the synthetic crRNA/tracrRNAs to a single cell suspension of dox treated iCas9 cells.

### Interactome analysis

Clarivate Analytics (previously Thomson Reuters) Computational Biology Methods for Drug Discovery (CBDD) toolkit implements several published algorithms (in R) for network and pathway analysis of –omics data. A proprietary R wrapper functioned to facilitate the use of the CBDD toolkit to run causal reasoning algorithms (Chindelevitch et al., 2012; Jaeger et al., 2014;). The knowledgebase used was a combination of a MetaBase (a manually curated proprietary database of mammalian biology from Clarivate Analytics) and STRING (Szklarczyk et al., 2015).

### OCT4 targeting assay

hPSCs were pre-treated with 1uM thiazovivin for at least 2 hours and harvested using accutase. A mixture of 4ug of Oct4-tdTomato-puroR targeting vector, 1ug of each gRNA cloned into a vector that co-expresses Cas9-D10A (or a vector lacking gRNAs as a control), and 2ug of either an episomal vector for p53DD (pCE-mP53DD) or EBNA1 alone (pCXB-EBNA1) were electroporated into 1×10^6 cells using a Neon electroporation system (Thermo). Cells were deposited into one well of a 6-well dish coated with matrigel containing 50% fresh mTESR:50% conditioned mTESR supplemented with bFGF (10ng/mL) and thiazovivin. After 48 hours, cells were selected with 0.3ug/mL puromycin in the presence of thiazovivin.

### Cas9/sgRNA ribonucleoprotein (RNP) complex delivery

1ul of 61uM Cas9 protein (IDT-1074182) was complexed with 1 ul of 100uM full length synthetic sgRNA (Synthego) for 5 minutes at room temperature. Following incubation RNP complexes were diluted with 100ul R buffer. Diluted RNPs were mixed with 1×10^6 pelleted hPSCs and electroporated at 1100v for 2 20ms pulses using the Neon electroporation system (Thermo) (Liang et. al., 2015).

## ACKNOWLEDGEMENTS

We thank Frederic Sigoillot for access to list of sgRNAs with multiple perfect binding sites. We thank Melody Morris and Abby Hill for help with the Interactome analysis. We thank Marc Hild for the constructive feedback on project.

## AUTHOR CONTRIBUTIONS

R.J.I. and A.K designed all experiments and wrote the manuscript. R.J.I. designed iCas9 constructs. R.J.I and S.K. made transgenic cell lines and characterized them. D.H. and C.Y. developed and performed indel analysis of mutated DNA samples. M.S. packaged the 47 individual lentiCRISPRs and K.A.W tested them. K.T. helped with live cell imaging of confluence. E.F., G.H. and G.M. helped with design of pooled screen, execution and analysis. J. R-H. generated sgRNA libraries. C.R. sequenced pooled screen samples. G.H., G.M., Z.Y., and W.F. provided access and analyzed non-targeting control data across transformed cell lines. T.K. identified sgRNAs with SNPs in H1-hESC genome. J.C. prepped RNA samples for RNA-seq experiments. R.R. performed RNA-seq. and interactome analysis. M.R.S. conducted high content image analysis. K.A.W. helped design and performed the *oct4* HDR assay

## SUPPLEMENTAL INFORMATION

**Figure S1.**
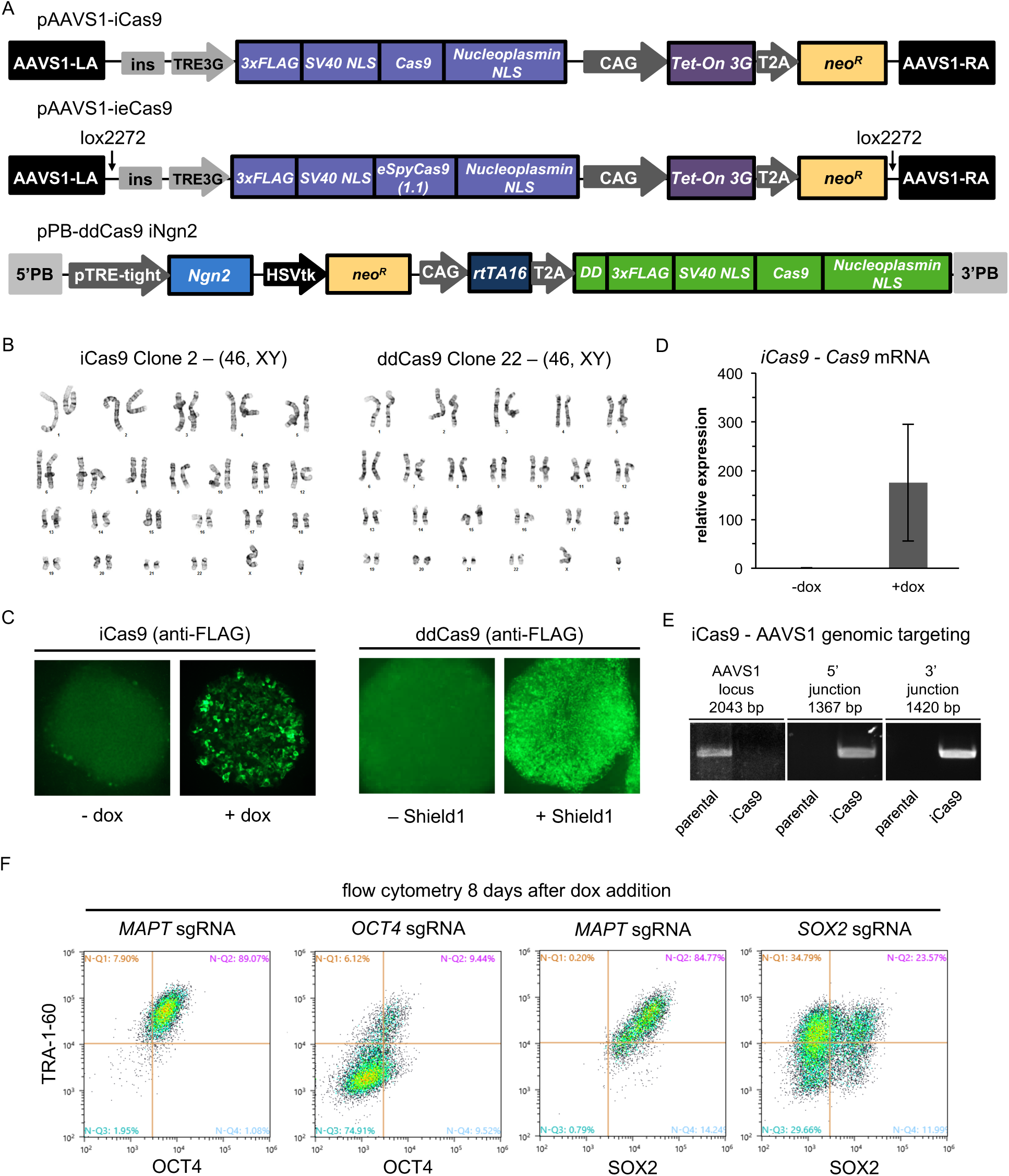
Inducible Cas9 constructs in hPSCs. (A) Detailed depiction of all-in-one dox inducible (pAAVS1-iCas9), inducible enhanced Cas9 (pAAVS1-ieCas9) and Shield1 inducible (pPB-ddCas9 iNgn2) Cas9 constructs. Although not utilized for this manuscript the ddCas9 transgene has an all-in-one dox inducible Ngn2 that can be used for rapid generation of cortical excitatory neurons from hPSCs. TRE3G and pTRE-tight, tetracycline response element promoter, ins, insulator CAG, constitutive promoter, Tet-On 3G and rtTA16, tetracycline transactivator protein, T2A, self-cleaving peptide, neo^R^, neomycin resistance gene, DD, destabilizing domain, PB, piggyBac repeats, LA, left homology arm, RA, right homology arm, HSVtk, herpes simplex virus (HSV) thymidine kinase promoter, ieCas9, dox inducible - enhanced Streptococcus Pyogenes Cas9 1.1 (eSpyCas9(1.1). (B) Karyotype analysis for the clones used in the study revealed no chromosomal abnormalities when the lines were first banked. 8402-iPSCs expressing ieCas9 are described by Wells et al., 2016 (C) Induction of Cas9 protein by addition of dox or Shield1 increase the amount of Cas9 protein detected by immunofluorescence using an antibody to detect FLAG tagged Cas9. (D) qPCR to detect *cas9* mRNA reveals that *cas9* expression is only induced in the presence of dox. Relative expression was calculated in comparison to untreated control. Bars represent mean expression. n=3 samples per condition. (E) Targeting of the iCas9 construct to the pAAVS1 safe harbor locus. Using a primer pair to span the AAVS1 knock-in site only amplifies in controls and indicates that iCas9 clone used in this study is homozygous. Junction PCR was used to detect both 5’ and 3’ specific junctions only in iCas9 transgenic cells. H1-ieCas9 is properly targeted homozygous as described by Wells et al., 2016. (F) iCas9 cells infected with *MAPT* targeting sgRNAs retain pluripotency marker expression. Control sgRNAs targeting OCT4 and SOX2 loose TRA-1-60 expression and either OCT4 or SOX2 expression respectively. Cells were fixed and stained after 8 days of dox treatment when on-target editing is near completion.

**Figure S2.**
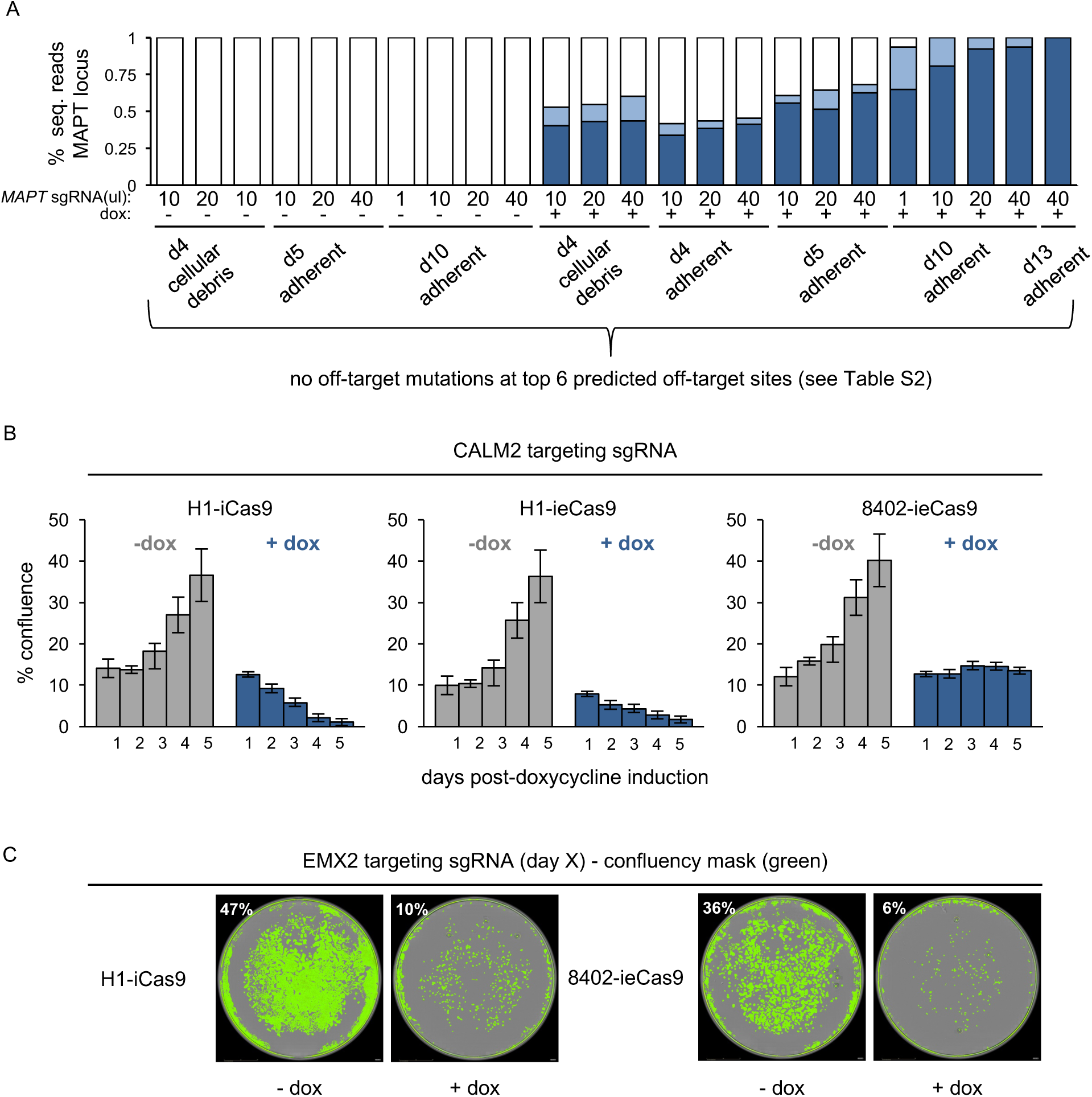
Cas9 toxicity is not related to off target activity. (A) On-target *mapt* indels in samples used for off-target analysis. Quantification of indels at *mapt* locus by NGS. Without dox, no indels are detected. With dox, frameshift and in-frame mutations increase over time. Cells were infected with 1-40ul lentivirus in 24-well plates seeded with 50,000 cells at the time of infection. Adherent samples were washed and dissociated while cellular debris was spun down from the spent media prior to dissociation and DNA isolation. Samples at day 5 and day 10 infected with 10ul of virus and treated +/− dox from figure 1D are displayed again for the context of off-target analysis. All samples were void of off-target mutations (Table S2). Control reads are represented by white bars, in-frame mutations by light blue bars and frameshift mutations by dark blue bars. (B) Quantification of toxic response to iCas9 and enhanced iCas9 induced DNA damage in H1-hESCs and 8402-iPSCs. Percent confluence was measured each day in cells expressing a *calm2* sgRNA grown with or without dox. Each bar represents mean confluence. Error bars +/− 1 std. dev from 16 images taken from 3 independent wells. (C) Whole well images from 24-well plates during editing with *emx2* sgRNA in H1-iCas9 and 8402-ieCas9 backgrounds. Both Cas9 and enhanced Cas9 variants cause toxicity in the presence of dox induced DSBs in hESCs or iPSCs. 3 wells per condition (n=3) average % confluence indicated top left. Confluency mask in green. ieCas9, dox inducible enhanced Streptococcus Pyogenes Cas9 1.1 (eSpyCas9(1.1)). H1, H1 human embryonic stem cells (hESCs), 8402, 8402 human induced pluripotent stem cells (iPSCs).

**Figure S3.**
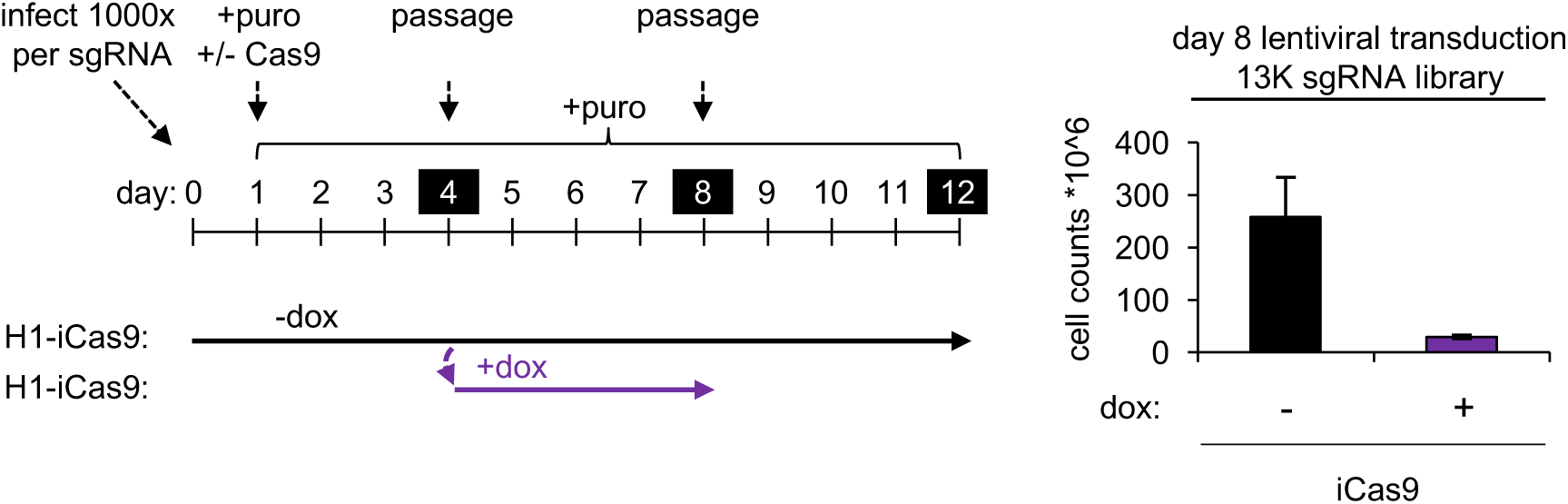
Dox treatment of infected sgRNA cell pool without previous exposure repeatedly reduces cell counts in four days. To repeat the large-scale toxic response to Cas9 activity the infected H1-iCas9 cells grown without dox were split into two conditions with or without dox. Four days following dox treatment the same pool of infected cells had a reduced cell count when cultured in the presence of both Cas9 and targeting sgRNAs. Bars represent average cell counts. Error bars +/− 1 std. dev from n=2 per condition.

**Figure S4.**
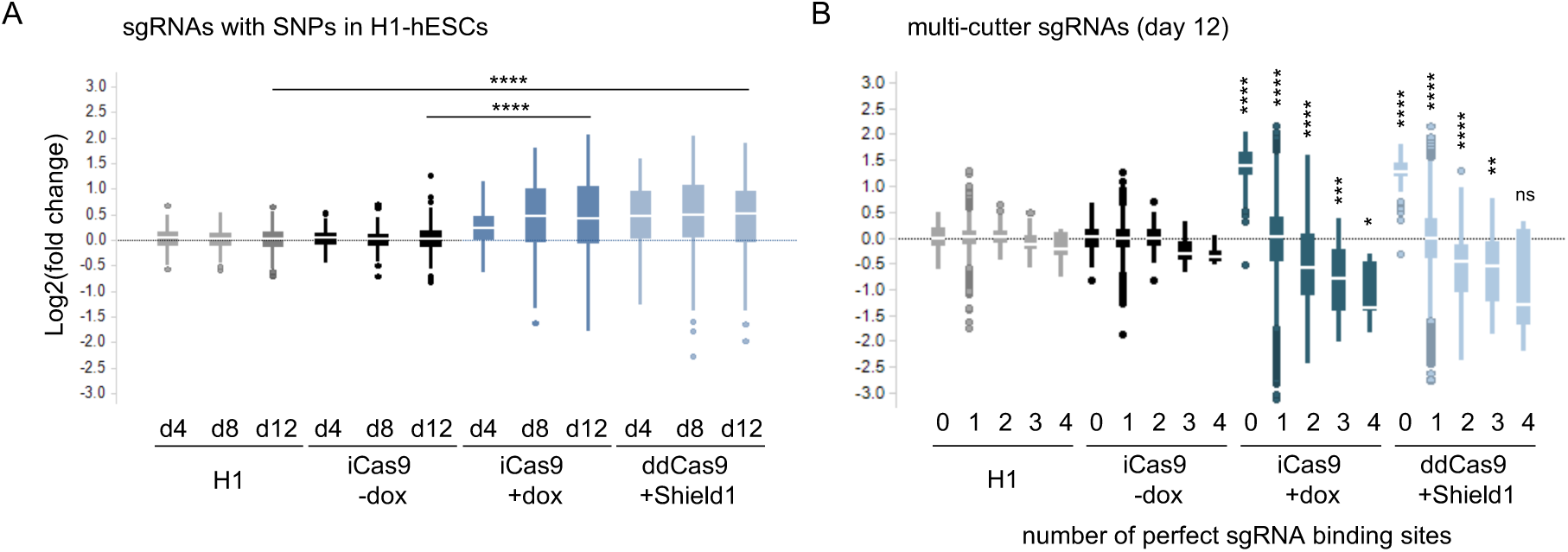
sgRNA design flaws are consistent with DSB toxicity in hPSCs. (A) log2(fold change) for 249 sgRNAs affected by SNPs in the H1-hESC genome. In the presence of Cas9, sgRNAs with binding sites disrupted by SNPs show an increase in representation. (B) log2(fold change) for 151 sgRNAs with one or more perfect cut sites. Only in the presence of Cas9, sgRNAs with no binding sites enrich while sgRNAs with 1 or more perfect binding sites dropout. Welch Two Sample t-test, unequal variance, Day 12, H1 compared to ddCas9, iCas9 minus dox compared to iCas9 plus dox, *p<0.05, **p<0.01, ***p<0.001, ****p<0.0001, n.s. – not significant. For each box blot the median is depicted by white line flanked by a rectangle spanning Q1-Q3.

**Figure S5.**
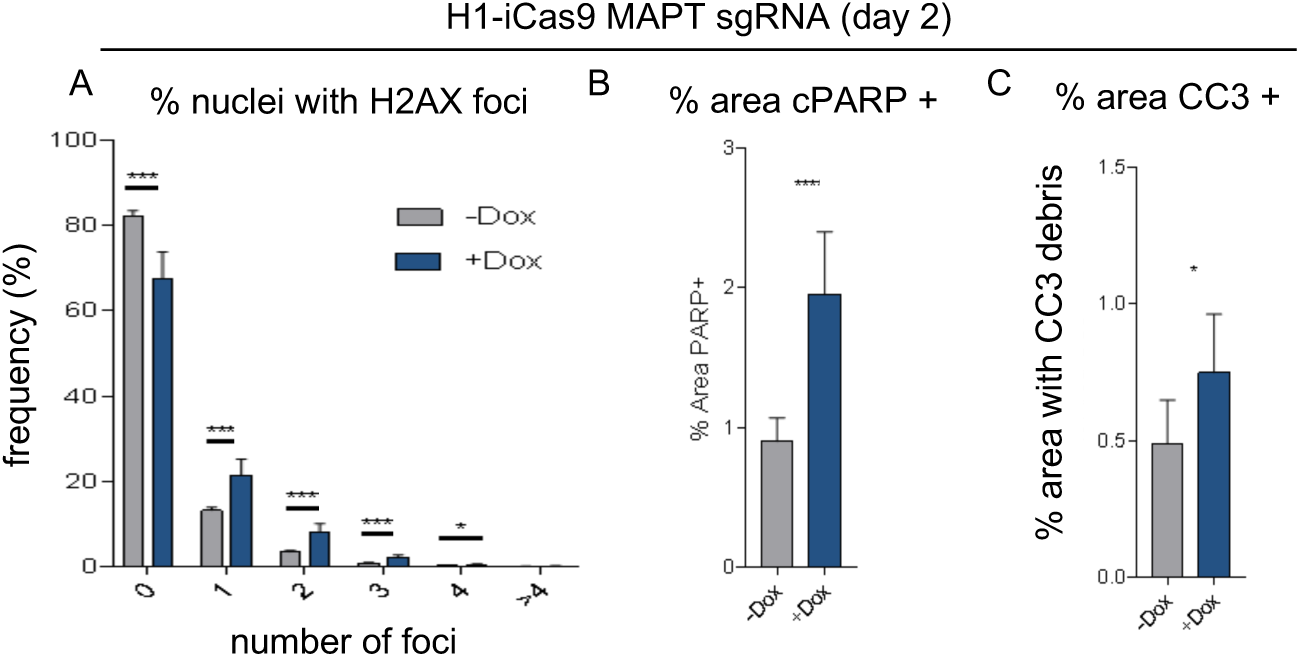
Cas9 induced DSBs increases DNA damage and cell death markers. High content image analysis of *mapt* sgRNA infected H1-iCas9 cells cultured with or without dox. (A) The frequency of nuclei containing pH2AX foci increases in the presence of a DSB. Unpaired Welch’s two-tailed t-test, n=8, unequal variance. (B) Percent area coverage of the apoptotic marker cPARP significantly increased in cells with a DSB. Unpaired Welch’s two-tailed t-test, n=8, unequal variance. (C) Immunostaining for cleaved caspase 3 (CC3), a marker for apoptosis, identified an increased area positive for CC3 debris within the DSB-induced (+dox) colonies. Unpaired Welch’s two-tailed t-test, n=8, similar variance. p<.05 ** p<.01 *** p<.001 **** p<0.0001. Bars represent average percent and error bars at +/− 1 standard deviation.

**Figure S6.**
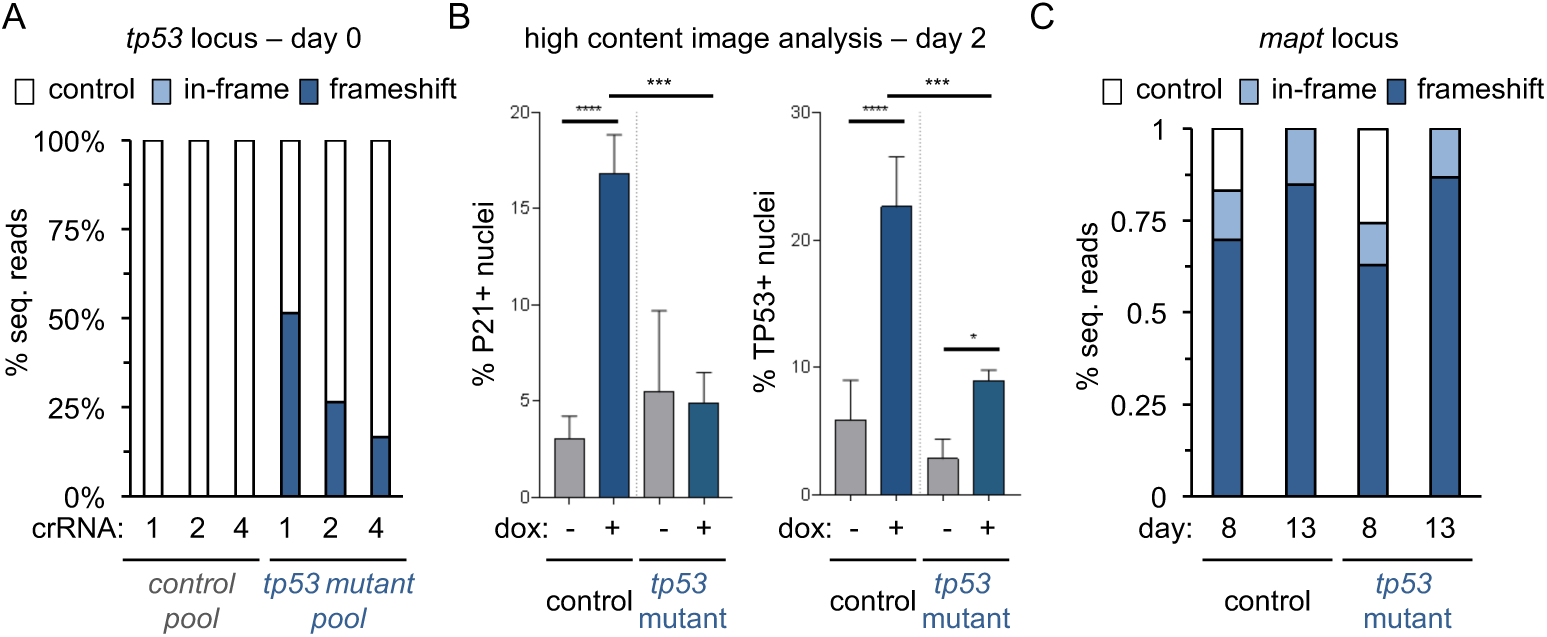
*tp53* mutant pool generation and analysis. (A) DNA from the onset of the experiment was isolated to quantify mutations at the *tp53* IOCUS by NGS. NO mutations are in the control pool while the mutant pool is a mix of wild-type and frameshift alleles at 3 different locations. NGS only measures one locus at a time and does not account for cis/trans mutations at other crRNA binding sites. Each mutation could be accompanied by either control reads or frameshift mutations at the other loci. The mutant pool therefore has a range from at least 50% and up to 93% *tp53* mutant alleles. Control reads are represented by white bars, in-frame mutations by light blue bars and frameshift mutations by dark blue bars. (n=1) (B) Quantification of DAPI stained nuclei positive for TP53 or P21 protein in control and *tp53* mutant pools infected with the *mapt* sgRNA cultured +/− dox for two days. Dox treated controls increase the percentage of TP53 or P21 positive nuclei, and this induction is significantly reduced in the *tp53* mutant pool. Error bars +/− 1 std. dev from n=4 wells. One-way ANOVA, similar variances. *p<.05, ***p<.001 **** p<.0001, n.s. – not significant. Bars represent average percent positive nuclei. (C) DNA isolated after 8- and 13-days of doxycycline treatment shows that on-target *mapt* editing efficiency is similar between control and *tp53* mutant pools. Average indels from three independent samples (n=3). Control reads are represented by white bars, in-frame mutations by light blue bars and frameshift mutations by dark blue bars.

**Figure S7.**
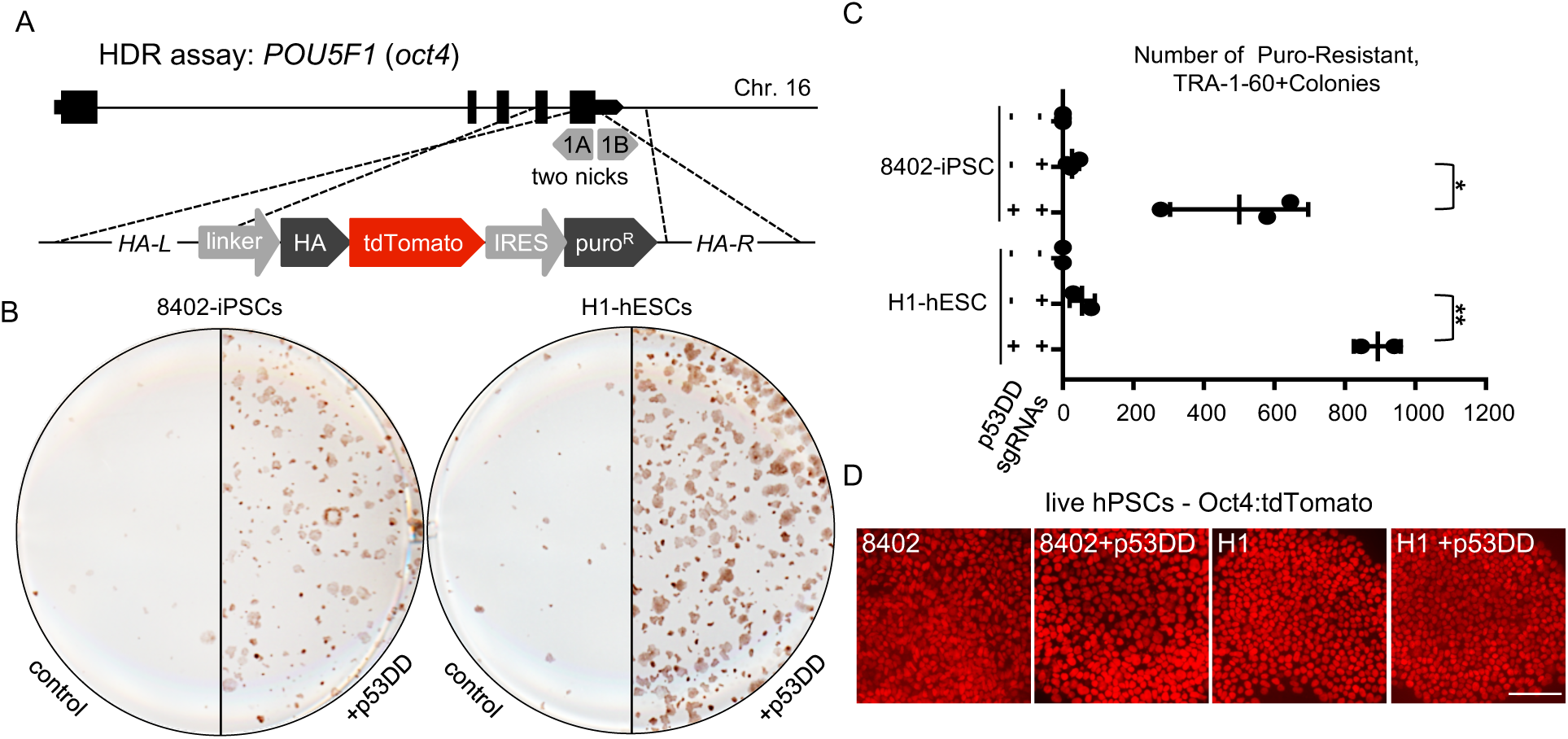
*tp53*-dependent toxicity inhibits Cas9 genome engineering in hPSCs. (A) Schematic of HDR assay targeting the *oct4/pou5f1* locus. A dual nickase targeting the stop codon was used to introduce a gene trap fusing an HA tagged tdTomato to the *oct4* ORF and an internal ribosome entry site (IRES) to drive the expression of the puro resistance gene off of the *oct4* promoter (B) TP53-induced toxicity inhibits the efficiency and yield of homology dependent repair (HDR) in hPSCs. Stem cell-specific TRA-1-60 antibodies conjugated to HRP were used to visualize colonies surviving puro selection following the electroporation of OCT4 donor, dual nickases and +/− p53DD plasmid. In H1-hESCs and 8402-iPSCs both the number and size of colonies with precise gene targeting are increased in the presence of p53DD relative to control. (C) Quantification of independent biological replicates conducted on different weeks in both 8402-iPSCs and H1-hESCs. unpaired, one-sided Welch’s t-test with unequal variance, *p<0.05, **p<0.01. 8402-iPSCs n=3, H1-hESCs is n=2‡. ‡Colonies were too large for accurate quantification in a 3^rd^ experiment. Mean represented by vertical center line (D) Live imaging of nuclear Oct4:tdTomato in both control and p53DD treated hPSCs. Scalebar = 100um.

**Table S1.**
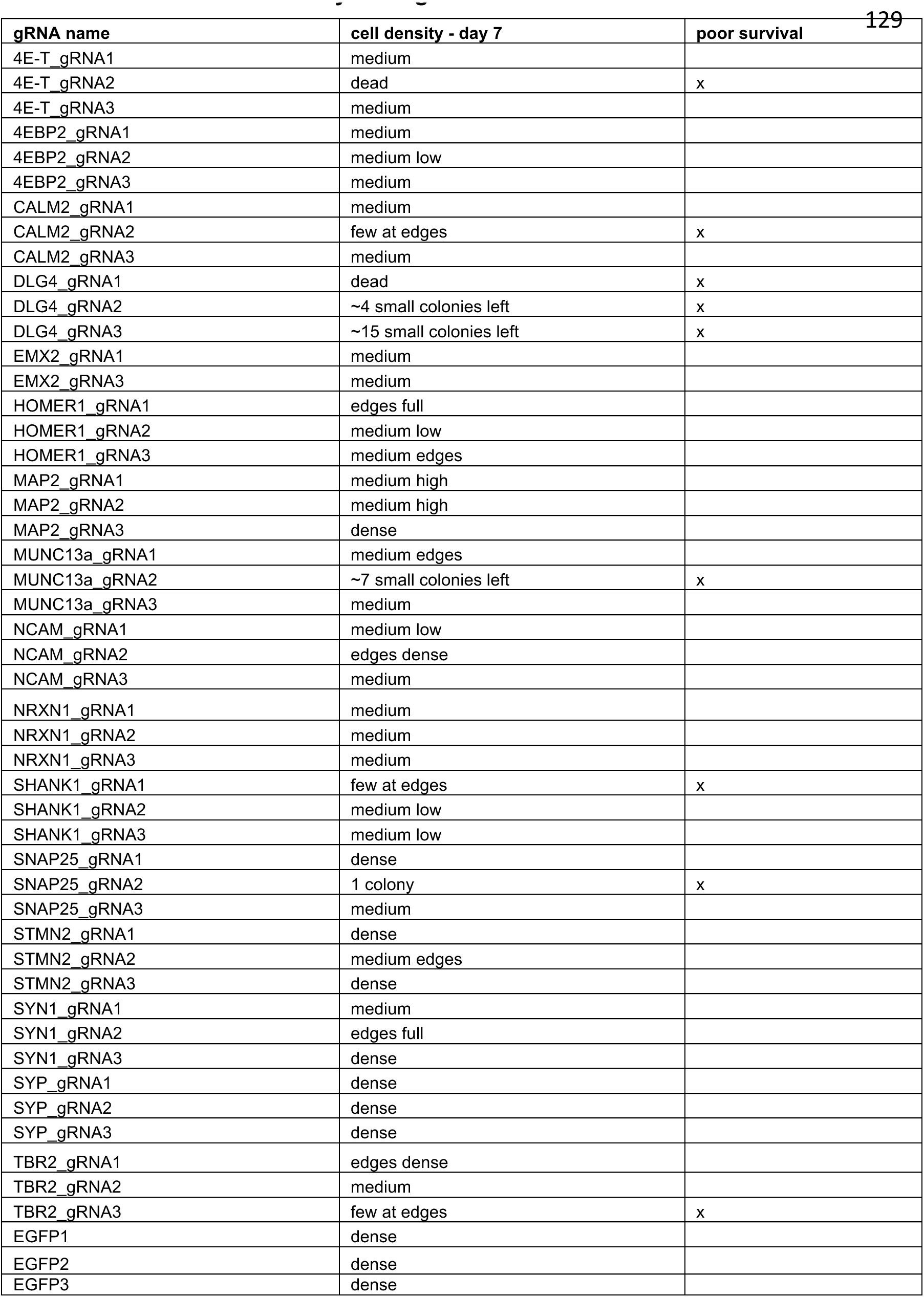
well to well variability - 47 sgRNAs.

**Table S2.**
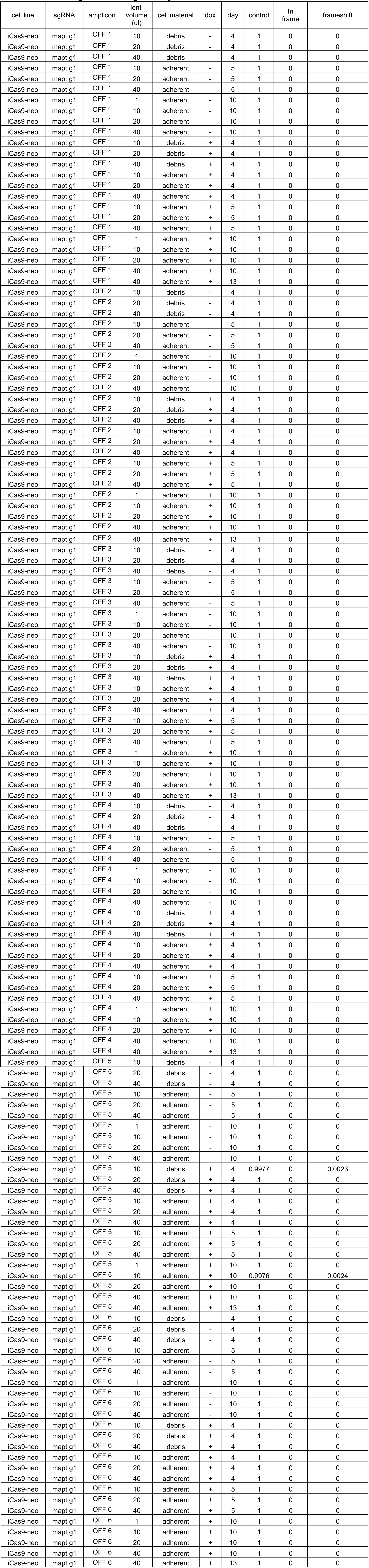
MAPT sgRNA off-target analysis.

**Table S3.**
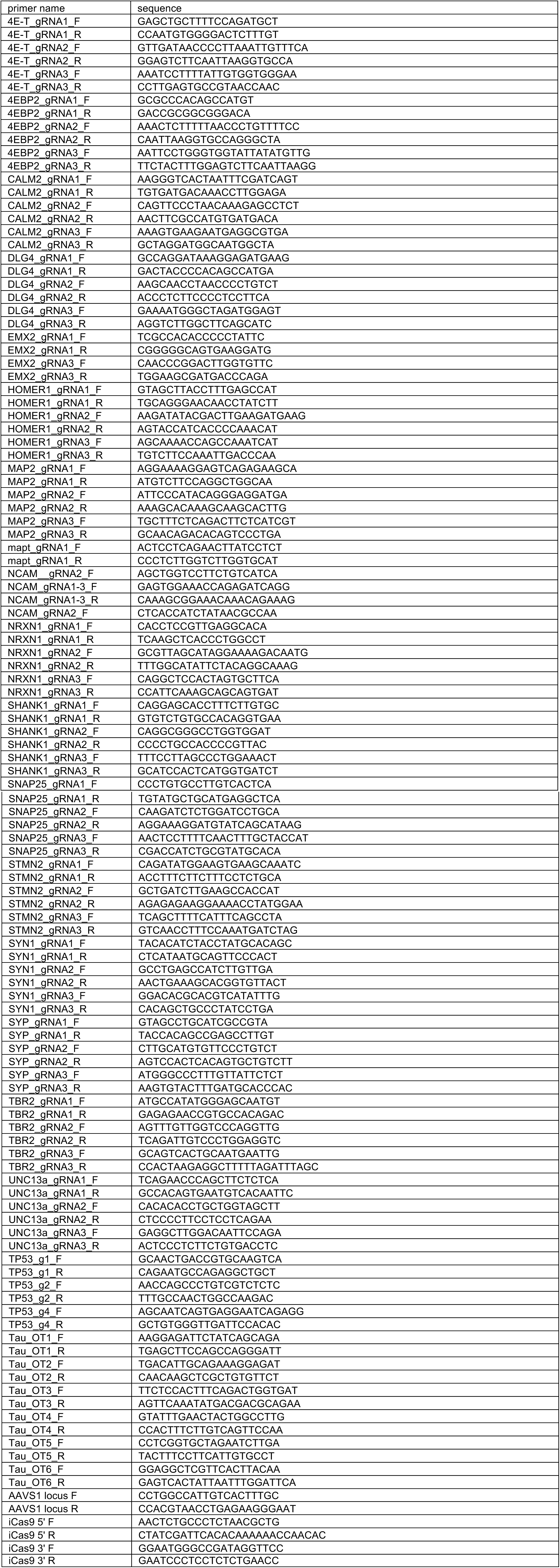
Primer sequences.

**Table S4.**
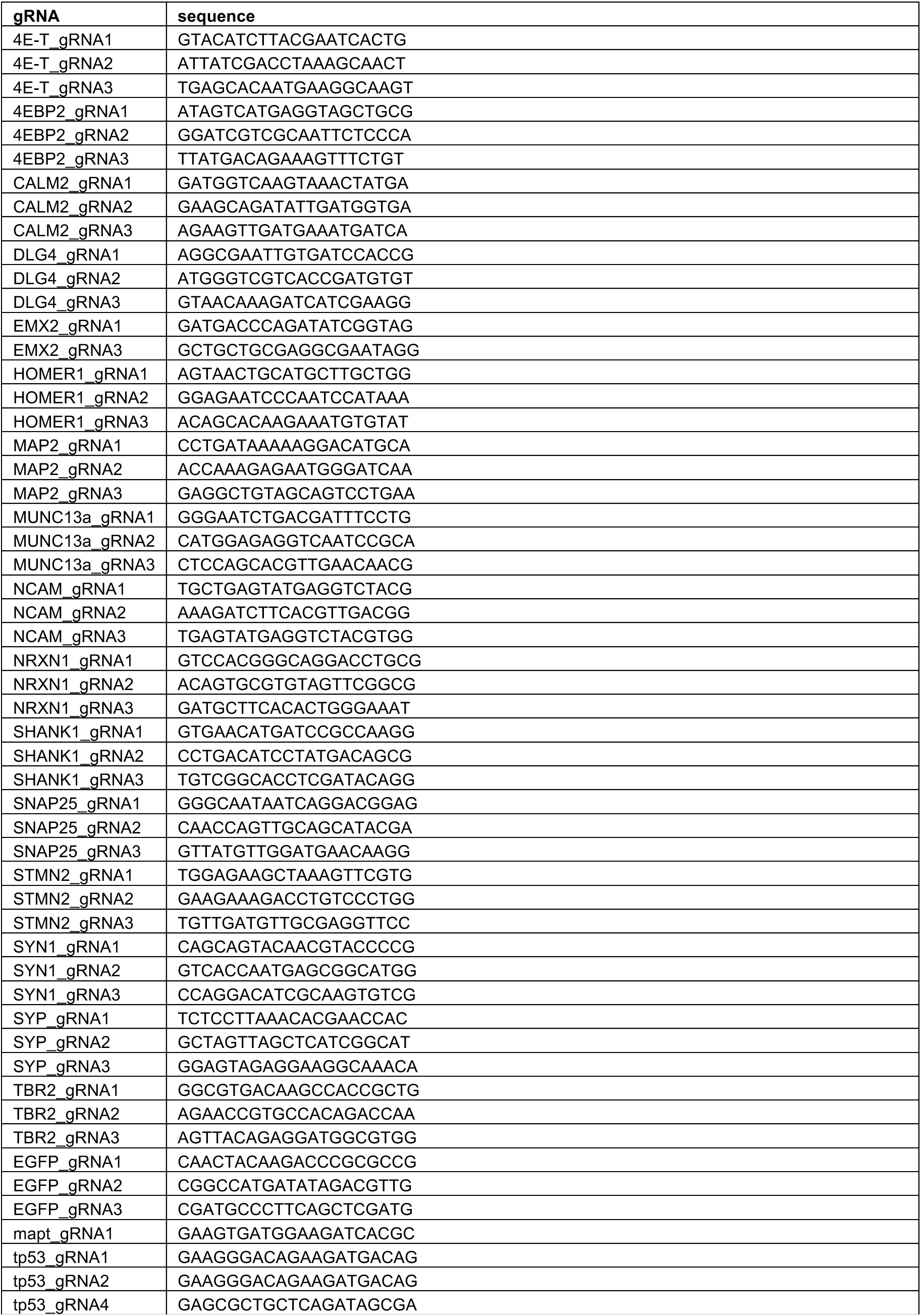
saRNA sequences.

### SUPPLEMENTAL EXPERIMENTAL PROCEDURES

#### hPSC cell culture

Cells were grown in TeSR-E8 media (STEMCELL TECH.-05940) on tissue-culture plates coated with vitronectin (Gibco-A14700). Passaging for maintenance was preformed using ReLeSR (STEMCELL TECH.-05873) to dissociate cell clumps to be replated in E8 plus thiazovivin (Selleckchem-S1459) at .2uM. For lentiviral transduction, electroporation, pooled screening and live imaging of confluence, accutase (Gibco-A1110501) was used to create a single cell suspension which was counted to accurately replate specific amounts of cells in E8 plus thiazovivin at .8uM. Karyotyping was performed by Cell Line Genetics (Madison, WI). H1-hESCs (WA01-NIHhESC-10-0043) were obtained from WiCell. 8402-iPSCs originated from GW08402 fibroblasts from the Coriell Institute and reprogrammed as described by Sun et al., 2016. hPSC lines were free of Myoplasma and tested using the Mycoalert Detection kit (Lonza). SNP fingerprinting confirmed the identify of hPSC lines used.

#### Inducible Cas9 cell line generation

Inducible Cas9 plasmids used in this study were synthesized by GenScript (Piscataway, NJ). Plasmid sequences available upon request. The AAVS1 TALE-Nuclease KIT(GE60xA-1) was obtained from System Biosciences (SBI). The iCas9 plasmid was knocked in to the AAVS1 locus of H1-hESCs, via electroporation using the NEON system (ThermoFisher). 1*10^6 cells with 1.5ug for each TALEN plasmid and 4ug for the donor plasmid were used for each electroporation at 1050V 30ms (2 pulses). After G418 selection, clones were picked, expanded and screened by treating with dox for 24 hours and staining for FLAG-tagged Cas9. Clones with strong expression of Cas9 were subsequently banked, karyotyped and were tested for proper targeting using junction PCR. The KOD Xtreme Hot Start DNA Polymerase (Millipore-71975) was used with the step-down cycling protocol as recommend by manufacturer for junction PCR. Primers are listed in Table S3. ddCas9 lines were electroporated using 4ug of the piggyBAC donor and 1ug of the piggyBAC dual helper plasmid. G418 resistant clones were selected by anti-FLAG stain and karyotyped. iCas9, and ddCas9 cell lines were maintained in media containing 200ug/ml G418 (Millipore-345812) to ensure proper transgene expression.

#### Lipid delivery of synthetic crRNA/tracrRNAs for Cas9 mutagenesis

iCas9 cells were treated with dox for 24h prior to transfection of synthetic crRNA/tracrRNA pairs. Cells were dissociated with accutase and replated at a density of 200,000 cells per well of 6-well plate in 2mL E8 plus thiazovivn. The amount of tracrRNA(90nM)/crRNA(30nM) used was calculated by referring to final concentration diluted 2mL of stem cell media (1 well of a 6-well plate). tracrRNAs/crRNAs were synthesized by IDT and resuspended in H20 at 100uM. tracrRNA (3X) and 3 crRNAs (1X each) targeting *tp53* were incubated together in H20 for 5 minutes at RT. The crRNA/tracrRNA mixture was then diluted in 100ul Opti-MEM (ThermoFisher-31985088) and incubated for 5 minutes at RT. In parallel 6ul of RNAimax (ThermoFisher-13778150) was diluted in 100ul Opti-MEM for 5 minutes at RT. Each tube was mixed and incubated for 15 minutes at RT. 200ul of the RNAimax/crRNA/tracrRNA/Opti-MEM was added dropwise to a well of a 6-well plate with freshly seeded iCas9 cells pretreated with dox. Cells were maintained in E8 media with doxycyline for 3 days following the transfection.

#### CRISPR indel analysis from genomic DNA

Template for PCR during library construction can be either cell lysate or genomic DNA purified using the Qiagen DNeasy Blood and Tissue Kit (Qiagen-69506) following the manufacturer’s protocol. For direct lysis, cells grown in a 96-well plate (Fig. 1B) were washed once with 100ul 1X PBS. Following removal of PBS, 40ul lysis buffer was added (10 mM Tris-HCl, 0.05% SDS, 25ug/ml proteinase K) to the cells, then incubated at 37C for an hour, followed by 85C proteinase inactivation for 15 minutes. The lysate was directly used as template for subsequent PCR. Each target was amplified using locus specific primers (Table S3) followed by a second amplification with illumina index containing primers. Libraries were quantified and sequenced on the Illumina MiSeq. For sequence analysis, raw reads were aligned to a reference sequence and binned into one of three categories – control, in-frame, frameshift.

#### LentiCRISPR transduction for Cas9 mutagenesis

The 47 sgRNAs in Figure 1 were designed using the sgRNA Designer (Broad Institute) (Table S4) and cloned into the pNGx_LV_g003_HA_Puro backbone by GenScript. The 13K lentiviral sgRNA library was designed, cloned into the pNGx_LV_g003_TagRFP_T2A_Puro backbone and packaged as described by Dejesus et al., 2016. For pooled screening viral titer was determined by exposing cells to a 12-point dose response of each lentiviral stock. 2*10^5^ cells were plated into a single well of a 6-well plate (2.1*10^4^ cells/cm^2^). Four days after infection % RFP was assayed by FACS (SONY SH800Z) and the data was used to calculate the amount of virus needed for .5 MOI. Puromycin concentration was optimized by infecting at .5 MOI and testing a dose-response of puro spanning .3ug/ml to 2ug/ml. At 2ug/ml puromycin 100% of surviving cells are RFP positive.

#### Pooled Screening

The 13,000 sub-genome library, included sgRNAs targeting 2.6K genes and non-targeting controls, was designed, synthesized, cloned and packaged as described by Dejesus et al., 2016. To infect at 1000x coverage 5 T225 flasks were seeded at 2.1*10^4^ cells/cm^2^ and infected at .5 MOI for each condition. Two independent replicates were maintained for each condition. 24h after infection cells were treated with .5ug/ml puromycin of for the remainder of the screen. Dox and Shield1 were added to the Cas9 positive conditions from day 1 through day 12. At each passage cells were counted to maintain 1000x coverage for both the newly seeded flask and the pellet for DNA isolation. To generate log2(fold change) values, DNA was isolated from pelleted cells and PCR amplified with primers targeting the lentiviral sgRNA backbone. Next generation library construction, sequencing and data analysis was performed as described by Dejesus et al., 2016. Non-hPSCs pooled screening data is available but restricted to non-targeting sgRNA sequences.

#### Live imaging of confluency

An IncuCyte zoom (Essen Biosciences) was used to quantify confluency in live cells each day post-media change. The confluence processing analysis tool (IncuCyte Zoom Software) calculated confluency for each sample. Average confluency and standard deviation was calculated for 96-well plates by taking a single image per well across multiple wells. Infected cells were allowed to recover after puro selection and were maintained in the absence of dox as to not induce mutations prior to the growth assay. For each population, cells were counted and plated in media containing dox at a density of 2.1*10^4^ cells/cm^2^ on 96-well plates at the start of the experiment (day 0).

#### RNA-seq, and qPCR

To detect signal from dying cells samples were collected by pelleting both the cellular debris in the media as well as the dissociated, formerly adherent, cells from an entire well per replicate in the same microcentrifuge tube. Total mRNA was isolated from using the RNeasy Mini kit plus (Qiagen-74134).

The Agilent 2100 bioanalyzer and the Nano 6000 kit (Agilent-5067-1511) were used to quantify and check the quality of each mRNA sample. 240ng of high quality RNA (RIN 10) was used for PolyA+ RNA-seq. Libraries were made using a Hamilton automated protocol with the TruSeq^®^ Stranded mRNA LT sample prep kit (Illumina-RS-122-2101) and sequenced on the Illumina HiSeq 2500. An average of more than 50 million 76-bp paired-end reads was obtained per sample. Processing was conducted using open source software. Raw fastq files were aligned to a human reference genome (GRCh37.74) using the STAR aligner (v2.5.1b) (Dobin et al., 2013). Gene counts and transcript quantification values (TPM) was performed using HTSeq-count (v0.6.0) (Anders et al., 2014) and RSEM (v1.2.28) (Li and Dewey, 2011) respectively. The gene counts were then used for differential expression analysis using DESeq2 (Love et al., 2014). 93% of variance is explained with principal component analysis and confirmed samples have similar variance.

For qPCR mRNA concentration was measured using a Nanodrop 2000 (Thermo Scientific). 200ug of RNA was used as template for cDNA synthesis using the SuperScript III firststrand synthesis system (ThermoFisher-18080051). cDNA was diluted 1:5 in H20 prior to analysis using taqman gene expression arrays and the 2x Fast Start Universal Probe master mix (ROX) (ROCHE-04913957001). 384-well qPCR plates were run on a ViiA 7 Real-Time PCR System (ThermoFisher). Relative expression was calculated as described by Pfaffl et al., 2002 and bACTIN was used as the reference gene. TaqMan gene expression arrays FAM-MGB (ThermoFisher-4331182); CDKN1A (Hs00355782_m1), bACTIN(Hs01060665_g1) fas (Hs00163653_m1). A custom TaqMan gene expression assay was ordered to detect Cas9 mRNA.

#### Immunofluorescence and Microscopy

Cells were fixed in 4% PFA in PBS for 10 minutes at room temperature and were washed with .1% triton X-100 in PBS after fixation. Cells were blocked in 2% goat serum, .01% BSA and .1% triton X-100 in PBS for 1hr at room temperature. Primary antibodies were diluted in blocking solution and incubated with cells over night at 4C. Cells were washed 3 times before incubation with secondary antibodies or fluorescently conjugated primary antibodies at room temp for 1.5 hours. Cell were washed 3 times and incubated with DAPI 1:1000 for 5 minutes at room temp before imaging. Primary antibodies: 1:250 P21 (12D1) (CST-2947), 1:250 P53 (7F5) (CST-2527), 1:300 FLAG (M2) (Sigma-F1804) 1:200 cleaved caspase-3 (Asp175) (CST-9661), 1:100 phospho-histone H2A.X (Ser139/Y142) (CST-5438), 1:50 Cleaved PARP-647 (Asp214) (D64E10) (CST-6987), Secondary antibodies: 1:500 Goat anti-Mouse IgG (H+L) AF488 (ThermoFisher-A-11029), 1:500 Goat anti-Rabbit IgG (H+L) AF488 conjugate (ThermoFisher-A-11008). For OCT4 targeting assay live cells were imaged for tdTomato fluorescence and then fixed, permeabilized, washed incubated with peroxidase suppressor (Thermo) for 30 min, washed twice, and then blocked for 30 min (5% goat serum/0.1% Tween-20/PBS). Cells were incubated at 37 degrees for 2 hours with anti-TRA-1-60 (MAB4360, Millipore, 1:300 dilution), washed 3 times, and then for 1 hour with anti-IgM conjugated to HRP (31440, Thermo, 1:250). A metal enhanced DAB substrate kit was used for detection (34065, Thermo). Live and fixed immunofluorescent images were taken using the 10x and 20x objectives on an Axio Observer.D1 (Ziess). Images for high content analysis were taken on an Incell 6000 (GE healthcare life sciences). TP53, P21, H2AX, cPARP, and CC3 immunoflurescence quantification was conducted via CellProfiler software. For TP53 and P21 proteins, average immunofluorescent intensity was determined for each nucleus, and a positive-expression threshold was set based on the nosecondary control. To quantify H2AX foci, the number of individual foci were detected within each nucleus via CellProfiler’s object detection module. To quantify cPARP and CC3, positive regions were detected via thresholding, and the area of this region was normalized to total plate area covered by colonies.

#### FACS

Cells were dissociated using accutase for 10 min at 37C to create a single cell suspension which was subsequently fixed in 4% PFA in PBS for 10 minutes at room temperature on a rocker. Cells were spun down at 300 RCF for 3 min between each subsequent solution change. Cells were washed with .1% Triton-X in PBS after fixation and blocked in 2% goat serum, .01% BSA and .1% triton X in PBS for 1hr at room temperature. Conjugated primary antibodies were diluted in blocking solution and incubated with cells on a rocker over night at 4C. 1:50 FITC conjugated anti-TRA-1-60 antibody (FCMAB115F – Millipore), 1:50 647 conjugated anti-OCT4 (C30A3) antibody (5263-CST), 1:50 647 conjugated anti-Sox2 (D6D9) antibody (5067-CST). Cells were washed and resuspended in PBS and transferred to a 5ml flow cytometry tube with strainer cap prior to FACS analysis on a SONY SH800Z. iCas9 ells were infected with lentiCRISPRs targeting *MAPT*, *OCT4*, and *SOX2* and were cultured fore 8 days in the presence of dox before FACS analysis. sgRNAs; OCT4-1 - CAACAATGAAAATCTTCAGG, SOX2-2 - CGTTCATCGACGAGGCTAAG.

